# Bifrost – Highly parallel construction and indexing of colored and compacted de Bruijn graphs

**DOI:** 10.1101/695338

**Authors:** Guillaume Holley, Páll Melsted

## Abstract

**Motivation:** De Bruijn graphs are the core data structure for a wide range of assemblers and genome analysis software processing High Throughput Sequencing datasets. For population genomic analysis, the colored de Bruijn graph is often used in order to take advantage of the massive sets of sequenced genomes available for each species. However, memory consumption of tools based on the de Bruijn graph is often prohibitive, due to the high number of vertices, edges or colors in the graph. In order to process large and complex genomes, most short-read assemblers based on the de Bruijn graph paradigm reduce the assembly complexity and memory usage by compacting first all maximal non-branching paths of the graph into single vertices. Yet, de Bruijn graph compaction is challenging as it requires the uncompacted de Bruijn graph to be available in memory.

**Results:** We present a new parallel and memory efficient algorithm enabling the direct construction of the compacted de Bruijn graph without producing the intermediate uncompacted de Bruijn graph. Bifrost features a broad range of functions such as sequence querying, storage of user data alongside vertices and graph editing that automatically preserve the compaction property. Bifrost makes full use of the dynamic index efficiency and proposes a graph coloring method efficiently mapping each *k*-mer of the graph to the set of genomes in which it occurs. Experimental results show that our algorithm is competitive with state-of-the-art de Bruijn graph compaction and coloring tools. Bifrost was able to build the colored and compacted de Bruijn graph of about 118,000 Salmonella genomes on a mid-class server in about 4 days using 103 GB of main memory.

**Availability:** https://github.com/pmelsted/bifrost available with a BSD-2 license

## 1 Introduction

The de Bruijn graph is an abstract data structure with a rich history in computational biology as a tool for genome assembly (Pevzner *et al.*, 2001; Idury and Waterman, 1995). With the advent of High Throughput Sequencing (HTS), the Overlap Layout Consensus (OLC) framework frequently used to assemble Sanger sequencing data (Yang *et al.*, 2011) was progressively replaced in favor of de Bruijn graph based methods. Since 2008, a wide range of genome assemblers based on the de Bruijn graph have been released (Chaisson and Pevzner, 2008; Zerbino and Birney, 2008; Simpson *et al.*, 2009; Luo *et al.*, 2012; Chikhi and Rizk, 2013; Bankevich *et al.*, 2012; MacCallum *et al.*, 2009). Although SMS (Single Molecule Sequencing) technologies (Rang *et al.*, 2018; Rhoads and Au, 2015) have re-introduced the OLC framework as the method of choice to assemble long and erroneous reads (Koren *et al.*, 2017; Li, 2016; Chin *et al.*, 2016; Kamath *et al.*, 2017), de Bruijn graph based methods are nonetheless used to assemble and correct long reads (Salmela and Rivals, 2014; Ruan and Li, 2019). Overall, the de Bruijn graphs have found widespread use for a variety of problems such as de novo transcriptome assembly (Robertson *et al.*, 2010), variant calling (Uricaru *et al.*, 2015), short read compression (Benoit *et al.*, 2015), short read correction (Limasset *et al.*, 2019), long read correction (Salmela and Rivals, 2014) and short read mapping (Liu *et al.*, 2016) to name a few. The colored de Bruijn graph is a variant of the de Bruijn graph which keeps track of the source of each vertex in the graph (Iqbal *et al.*, 2012). The initial application was for assembly and genotyping but it has also found use in pan-genomics (Zekic *et al.*, 2018), variant calling (Fang *et al.*, 2016) and transcript quantification methods (Bray *et al.*, 2016).

Despite serving as a building block for many methods in computational biology, the de Bruijn graph adoption is hindered by two factors. First, the memory usage and computational requirements for building de Bruijn graphs from raw sequencing reads are considerable compared to alignment to a reference genome while only a handful of tools have focused on de Bruijn graph compaction (Minkin *et al.*, 2016; Chikhi *et al.*, 2016; Marcus *et al.*, 2014; Baier *et al.*, 2016; Minkin *et al.*, 2013). Second, de Bruijn graph construction usually requires tight integration with the code. In the best case, software libraries for building and manipulating de Bruijn graphs are used (Drezen *et al.*, 2014; Crusoe *et al.*, 2015) but in most cases, data structures to index the de Bruijn graph are re-implemented. Those downsides are intensified in the colored de Bruijn graph for which the memory consumption of colors rapidly overtakes the vertices and edges memory usage (Almodaresi *et al.*, 2017). For this reason, a lot of attention has been given to succinct data structures for building the colored de Bruijn graph (Marcus *et al.*, 2014; Holt and McMillan, 2014; Holley *et al.*, 2015; Baier *et al.*, 2016; Muggli *et al.*, 2017; Almodaresi *et al.*, 2017, 2018; Muggli *et al.*, 2019) and data structures for multi-set *k*-mer indexing (Solomon and Kingsford, 2016; Sun *et al.*, 2018; Solomon and Kingsford, 2018; Pandey *et al.*, 2018; Yu *et al.*, 2018; Bradley *et al.*, 2019).

In this paper, we present Bifrost, a software for efficiently constructing the colored and compacted de Bruijn graph, both in terms of runtime and memory usage. The data structures and algorithms implemented in Bifrost are specifically tailored for fast and lightweight construction, querying and dynamic manipulation of compacted de Bruijn graphs, both regular and colored. The software is designed to take advantage of multiple cores and modern processors instruction sets (SIMD operations). Bifrost is also available as a C++11 software library with minimal external dependencies and allows developers to build on top of an efficient de Bruijn graph engine by using the Bifrost API.

## 2 Definitions

A string *s* is a sequence of symbols drawn from an alphabet 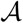. The length of *s* is denoted by *|s|*. A substring of *s* is a string occurring in *s*: it has a starting position *i*, a length *l* and is denoted by *s*(*i, l*). A substring of length *l* is also denoted an *l*-mer. In the following, we assume 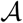 is the DNA alphabet 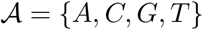 for which symbols have complements: (*A, T*) and (*C, G*) are the complementing pairs. The reverse-complemented string 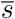 is the reverse sequence of complemented symbols in *s*. The canonical string 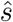 is the lexicographically smallest of *s* and its reverse-complement 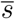. The minimizer (Roberts *et al.*, 2004; Grabowski *et al.*, 2015) of an *l*-mer *x* is a *g*-mer *y* occuring in *x* such that *g < l* and *y* is the lexicographically smallest of all the *g*-mers in *x*. The lexicographical order can be cumbersome to use since poly-A *g*-mers naturally occurs in sequencing data and is often replaced by a random order. The simplest way to obtain a random order is to compute a hash-value for each *g*-mer in *x* and select the *g*-mer with the smallest hash-value as the minimizer. In this work, we will only consider minimizers generated by random orderings.

A de Bruijn graph (dBG) is a directed graph *G* = (*V, E*) in which each vertex *υ ∈ V* represents a *k*-mer. A directed edge *e ∈ E* from vertex *υ* to vertex *υ*′ representing *k*-mers *x* and *x*′, respectively, exists if and only if *x*(2*, k −* 1) = *x*′(1*, k −* 1). Each *k*-mer *x* has 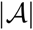 possible successors *x*(2*, k −* 1) ⊙ a and 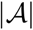 possible predecessors a ⊙ x(1*, k −* 1) in *G* with 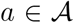 and ⊙ as the concatenation operator. Note that in the original combinatorial definition of the dBG, all possible *k*-mers for an alphabet 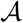 are present in the graph, whereas in computational biology, the definition is restricted to a subset of the de Bruijn graph representing the *k*-mers in the input. A path in the graph is a sequence of distinct and connected vertices *p* = (*υ*_1_, …, *υ*_m_). We say that the path *p* is *non-branching* if all its vertices have an in- and out-degree of one with exception of the head vertex *υ*_1_ which can have more than one incoming edge and the tail vertex *υ*_m_ which can have more than one outgoing edge. A non-branching path is maximal if he cannot be extended in the graph without being branching. A compacted de Bruijn graph (cdBG) merges all maximal non-branching paths of *η* vertices from the dBG into single vertices, called unitigs, representing words of length *k* + *η −* 1. Minimal examples of dBG and cdBG are provided in Figures 1a and 1b respectively. A colored de Bruijn graph is a graph *G* = (*V, E, C*) in which (*V, E*) is a dBG and *C* is a set of colors such that each vertex *υ ∈ V* maps to a subset of *C*, we extend the definition of a cdBG to a colored compacted de Bruijn Graph (ccdBG) to be a graph *G* = (*V, E, C*), where (*V, E*) is a cDBG, so the vertices represent unitigs, and each *k*-mer of a unitig maps to a subset of *C*.

**Figure 1:**
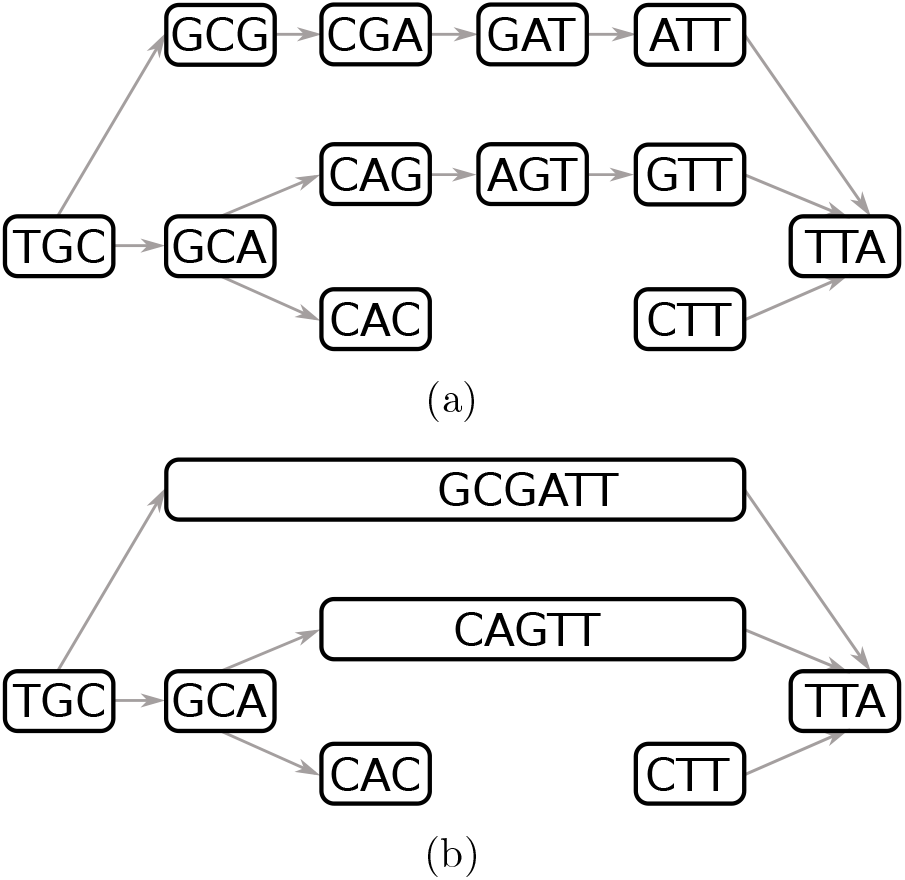
A de Bruijn graph in (a) and its compacted counterpart in (b) using 3-mers. For simplicity, reverse-complements are not considered.

Introduced by Bloom (1970), the Bloom filter (BF) is a space and time efficient data structure that records the approximate membership of elements in a set. The BF is represented as a bitmap *B* of *m* bits initialized with 0s, coupled with a set of *f* hash functions *h*_1_, …, *h*_f_. Inserting and querying an element *e* into *B* is performed with the functions

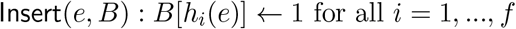

and

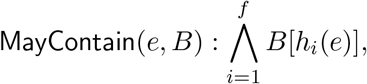

respectively, in which Λ is the logical conjunction operator. Those functions require 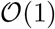 time. The function `MayContain` may report false positives when querying for elements which were never inserted but are present in *B* as a result of independent insertions. Given *n* elements to insert, the optimal number of hash functions to use (Kirsch and Mitzenmacher, 2006) is 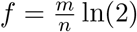, for an approximate false positive rate of

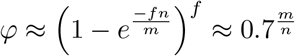

Hence, the BF trades off memory usage and time complexity with a decreased false positive rate.

In order to accelerate BFs, (Kirsch and Mitzenmacher, 2006) demonstrated that two hash functions combined in a double hashing technique can be applied in order to simulate more than two hash functions and obtain similar hashing performance. One main drawback of BFs is their poor data locality as bits corresponding to one element are scattered over *B*, resulting in several CPU cache-misses when inserting and querying. This issue was addressed in (Putze *et al.*, 2009), which presented the Blocked Bloom Filter (BBF), an array of smaller BFs individually fitting into one or multiple cache lines. To insert or look-up an element, a supplementary hash function is used to determine which BF to load. While BBFs are fast, their false positives ratios are usually higher than regular BFs due to the unbalanced load of each BF in the array.

As minimizers are used extensively throughout Bifrost, we use an efficient rolling hash function based on the work of Lemire and Kaser (2010) to select a *g*-mer as the minimizer within a single *k*-mer. Since overlapping *k*-mers are likely to share minimizers, we use an ascending minima approach (Harter, 2009) to recompute minimizers with amortized *O*(1) costs, so that iterating over minimizers of adjacent *k*-mers in a sequence is linear in the length of the sequence. Another optimization is to restrict the computation of minimizers to a subset of *g*-mers of a *k*-mer; namely, we exclude the first and last *g*-mer as a candidate for being a minimizer. This ensures that for a given *k*-mer, all of its forward, respectively backward, adjacent *k*-mers necessarily share the same minimizer. While it is likely that a *k*-mer *x* and its neighbor *x*′ share a minimizer, this neighbor hashing trick (Holley *et al.*, 2015) guarantees that when searching all forward, respectively backward, neighbors of *x*, they will all have the same minimizer and will be stored within the same block of a BBF, thus minimizing cache misses.

## 3 Methods

To build the cdBG, Bifrost builds first an approximation of the dBG using BBFs to filter out sequencing errors from the reads. The BBF containing the *k*-mers to insert in the graph is then used to build the exact cdBG. Finally, *k*-mers contained in the unitigs are mapped to their colors representing the input sources in which they occur.

The algorithms developed are described in this section. First Section 3.1 describes how an approximation of the true dBG is built from a set of sequencing reads, Algorithm 1 shows how to remove the majority of *k*-mers occuring only once and Algorithm 2 details the construction of a multithreaded BBF. Section 3.2 shows how the approximate cdBG is built from the BBF, Algorithm 3 details how the cdBG data structure indexes unitigs and Algorithm 4 shows how the data structure is queried for *k*-mers. Algorithm 5 shows how the unitigs of the approximate cdBG are discovered and Algorithm 6 constructs the approximated cdBG from a set of reads. Finally the approximate cdBG is cleaned up and converted to an exact cdBG using Algorithm 7. Section 3.3 shows how a ccdBG can be built efficiently on top of a cdBG with Algorithm 8.

### 3.1 Approximating the de Bruijn graph

The *k*-mers extracted from the reads will be inserted into two BBFs: *BBF*_1_ will contain all *k*-mers occurring at least once in the input read sets while *BBF*_2_ will contain all *k*-mers occurring twice or more often. This separation allows us to filter out unique *k*-mers which are likely to be sequencing errors (Melsted and Pritchard, 2011). Algorithm 1 starts by iterating over the reads and extracts all the canonical *k*-mers. *BBF*_1_ is queried for the presence of each such *k*-mer and *k*-mers already present in *BBF*_1_ are inserted into *BBF*_2_. Finally, *BBF*_1_ is discarded as the cdBG will be built from the *k*-mers of *BBF*_2_.

In order to accelerate the insertions into the BBFs, the minimizer hash-value of each *k*-mer is used to determine the BBF block in which the *k*-mer is inserted. This guarantees that overlapping *k*-mers sharing the same minimizer position within a read are inserted into the same BBF block, thus improving the cache efficiency of BBFs. Furthermore, the neighbor hashing of the minimizers guarantees that all predecessors and successors of a *k*-mer are hashing to the same block, thus improving graph traversal for the exact cdBG construction step. Finally, the BBFs in Bifrost use 2-choice hashing (Azar *et al.*, 1999) to balance the number of insertions per block and reduce the number of false positives.

#### Algorithm 1 Construct Blocked Bloom Filters

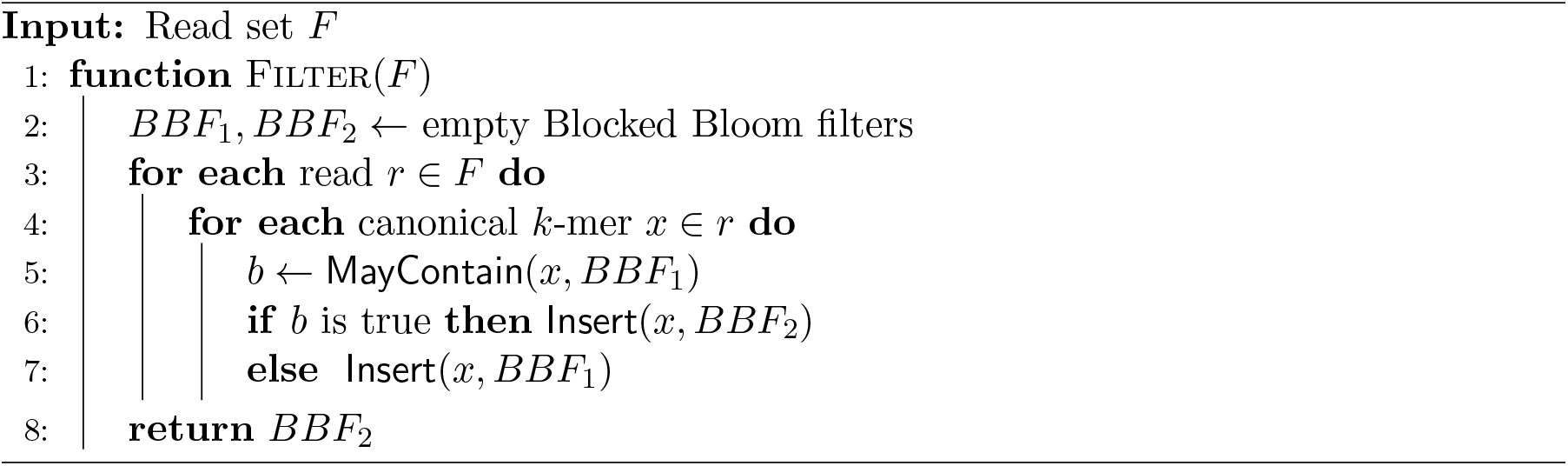

Instead of selecting a single BBF block when inserting a *k*-mer, two blocks are selected. If none of the two blocks already contains the *k*-mer, it is inserted into the block which has the fewest number of bits set. To enable parallel insertions, each BBF block is equipped with a spinlock to avoid multiple threads inserting at the same time within the same block. Algorithm 2 refines the insertion function introduced in Section 2 to enable 2-choice hashing and spinlocks usage with BBFs. Bifrost can make use of modern processors instruction sets to query simultaneously up to 16 bits within a block using AVX instructions.

#### Algorithm 2 Insert a *k*-mer into a BBF

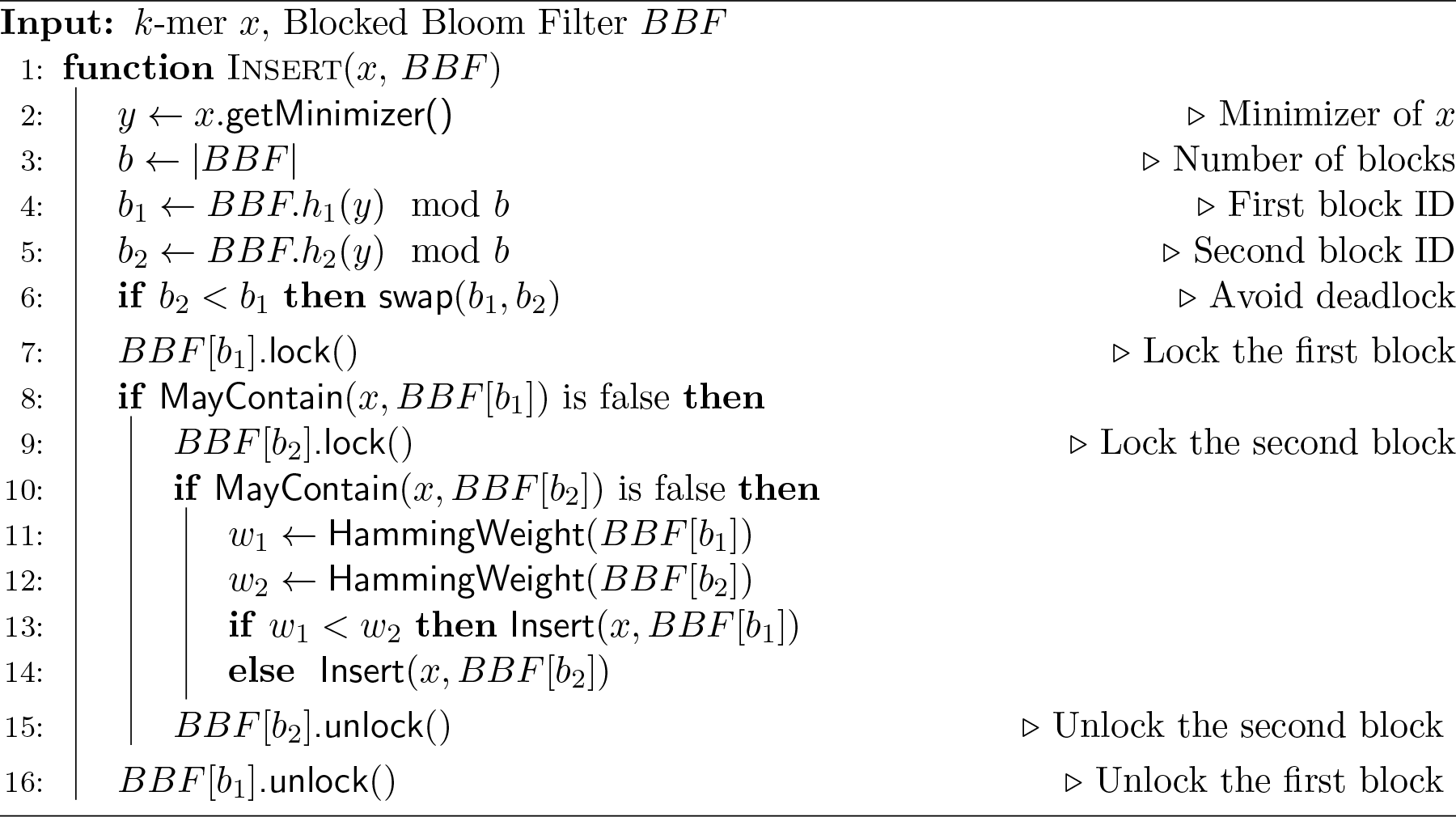

### 3.2 Constructing the compacted de Bruijn graph

The following section describes the data structure indexing the unitigs. Section 3.2.2 details the unitig extraction procedure from the BBF and the insertion of unitigs into the cdBG data structure.

#### 3.2.1 Data structure

The cdBG data structure *D* = (*U, M*) is composed of a unitig array *U* and a hash-table of minimizers *M*. A unitig *u* is first inserted into *U* and gets a unique identifier *id*_*u*_. Unitig *u* is then decomposed into its set of constituent *k*-mers from which minimizers are extracted. Each minimizer is identified by a position *p*_*m*_ in *u*. While there can be as many minimizer positions as there are *k*-mers in the unitig, it is likely that multiple overlapping *k*-mers share the same minimizer position. The canonical *g*-mers corresponding to the minimizers are inserted into *M* and associated with their position *p*_*m*_ in *u* and the identifier *id*_*u*_. Note that a minimizer might have multiple occurrences, either within a unitig or in different unitigs of the graph. The cdBG data structure *D* is illustrated in Figure 2. Algorithm 3 details the insertion of a unitig *u* in the cdBG data structure. Note that removing a unitig from the graph can be done in a reversed-fashion to Algorithm 3: The tuples associated with unitig *u* are removed from *M* and unitig *u* is removed from *U*.

**Figure 2:**
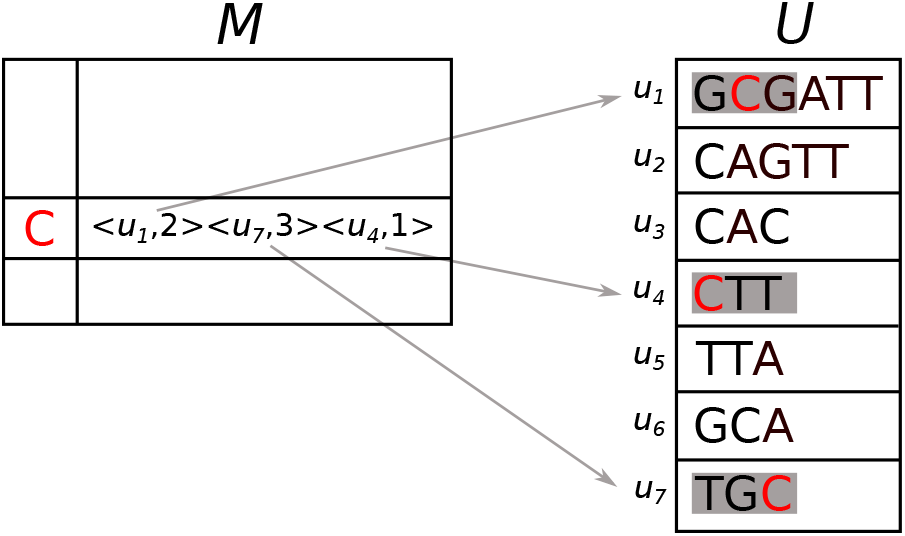
Data structure of a cdBG composed of a hash-table *M* and a unitig array *U*. Unitigs are composed of 3-mers and are indexed using minimizers of length 1. For simplicity, a lexicographic ordering of minimizers is here used and only one minimizer is shown.

##### Algorithm 3 Insert a Unitig into a cdBG

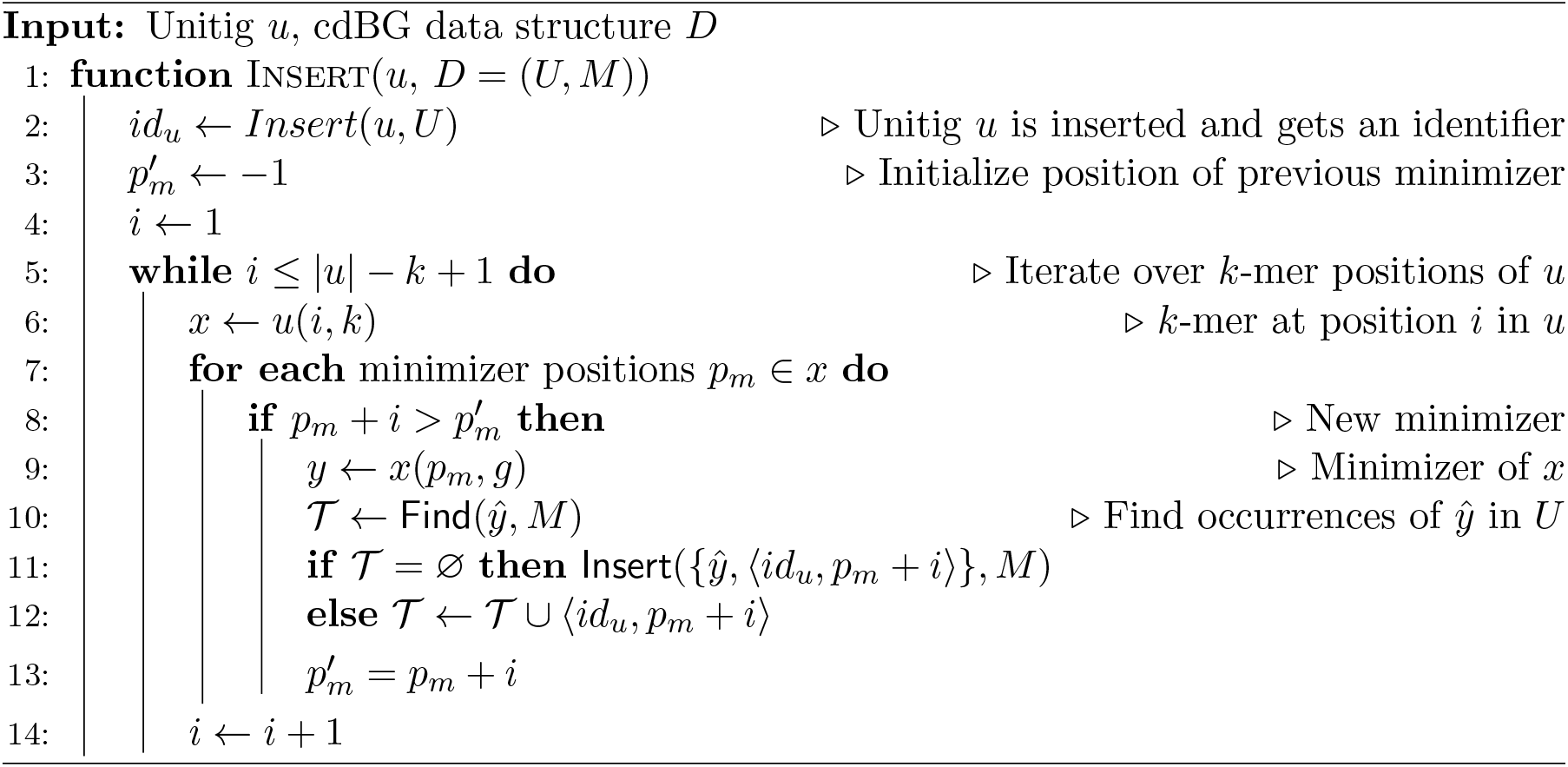

Looking-up a *k*-mer *x* in the cdBG data structure is similar to inserting a unitig. The canonical *g*-mer corresponding to the minimizer of *x* is extracted and used to query *M*. If the *g*-mer is not in *M*, *x* does not occur in a unitig of the cdBG. However, if the *g*-mer is present, the identifiers of the unitigs containing the *g*-mer and the *g*-mer positions within those unitigs are returned. K-mer *x* and its reverse-complement 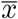 are then anchored in those unitigs at the given minimizer positions and compared. If the comparison is positive, a tuple with the unitig identifier and the *k*-mer position in the unitig is returned. Algorithm 4 shows how to look-up *D* for a *k*-mer.

##### Algorithm 4 Find a *k*-mer in a Unitig in a cdBG

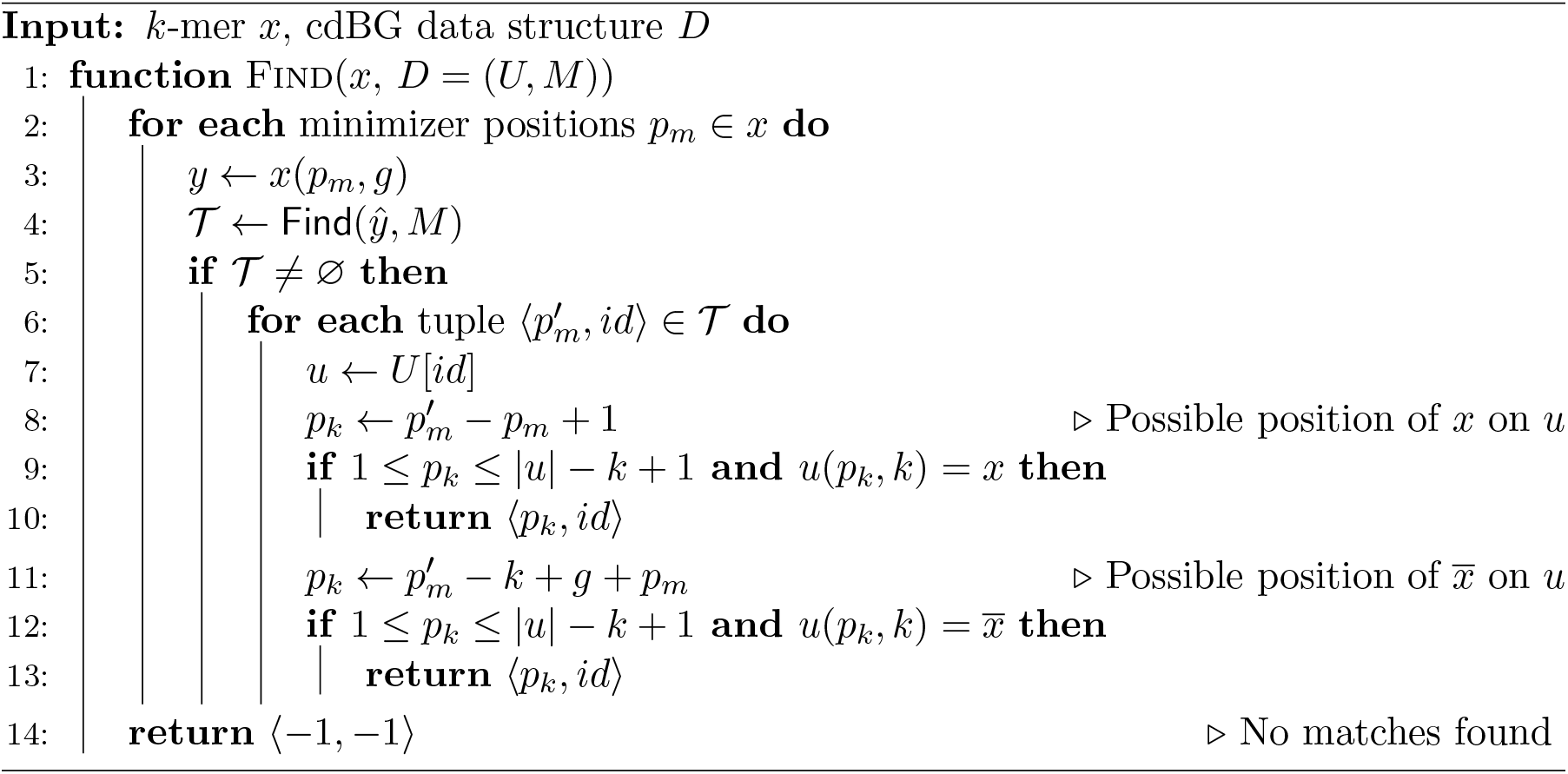

#### 3.2.2 Unitig extraction

The BBF returned by Algorithm 1 represents an approximation of the dBG: It contains the true positive *k*-mers composing the unitigs of the cdBG and false positive *k*-mers to exclude from the cdBG construction. The false positives *k*-mers are either artifacts of *BBF*_2_ or single occurrence *k*-mers that should have been filtered out by Algorithm 1 but were inserted into *BBF*_2_ as a result of their false occurrences in *BBF*_1_. Although BBFs are efficient data structures, they do not allow to iterate over the contents. To get around this limitation, we iterate over the original set of reads and query *BBF*_2_ to identify *k*-mers that are present.

Given a *k*-mer *x*, Algorithm 5 extracts from the BBF the unitig from which *x* is a substring, conditioned upon the presence of *x* in the BBF. K-mer *x* is extended forward, respectively backward, by reconstructing iteratively the prefix, respectively suffix, of the unitig using function `Extend`. Note that a backward extension is performed by extending forward from the reverse-complement of *x* and the extracted suffix is reverse-complemented to obtain the unitig prefix. Forward extensions are made with function `ExtendForward` which iteratively concatenate the last character from the next *k*-mer in the extension until no more *k*-mer is found or the extracted *k*-mer creates a cycle. Finally, *k*-mer *x* is extended with *x*′ using function `ExtendKmer` if the two *k*-mers belong to the same maximal non-branching path, i.e, if *x*′ is the only successor of *x* in the BBF and *x* is the only predecessor of *x*′ in the BBF,

##### Algorithm 5 Unitig extraction from a BBF

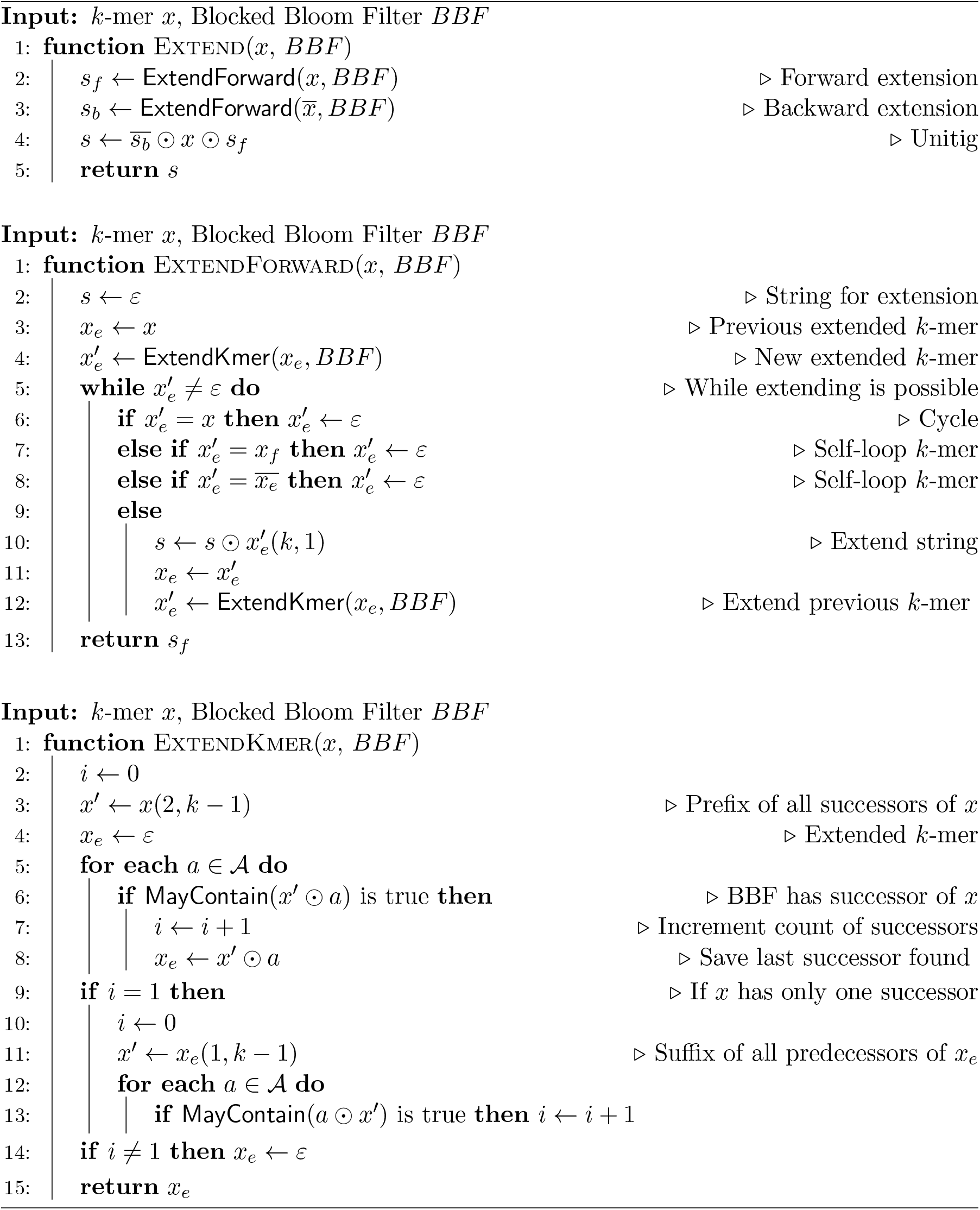

Given the read set, the BBF containing the filtered *k*-mers and an empty cdBG data structure, Algorithm 6 extracts the unitigs from the BBF and inserts them into the cdBG data structure. The algorithm iterates over the *k*-mers of the reads and query the BBF for their presence. A missing *k*-mer in the BBF indicates the *k*-mer was filtered out by Algorithm 1 and will not be part of a unitig, in which case the next *k*-mer in the read is queried. However, in case of the *k*-mer presence in the BBF, the cdBG is searched for the unitig containing this *k*-mer using Algorithm 4. If the *k*-mer is missing from the unitigs present in the cdBG data structure, it means its unitig has not been extracted yet from the BBF. The extraction using Algorithm 5 takes place and the extracted unitig is inserted into the cdBG data structure with Algorithm 3.

##### Algorithm 6 Initial cdBG Construction

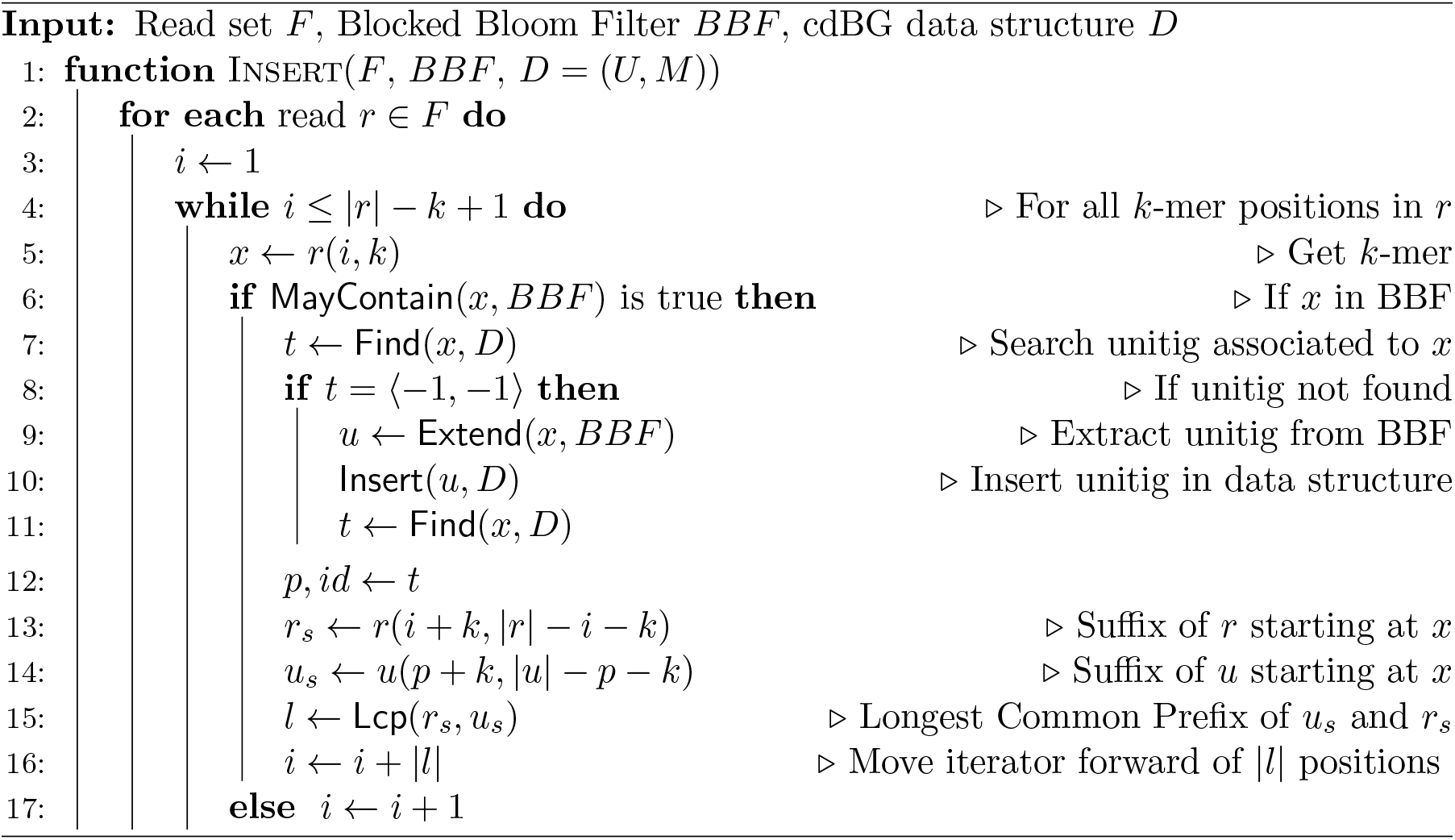

#### 3.2.3 Eliminating the false positive*k*-mers

The cdBG constructed by Algorithm 6 is not exact as it contains false positive *k*-mers of *BBF*_2_. Those false positive *k*-mers create two types of errors in the graph:

- False connection: A false positive *k*-mer connects a unitig with no successors to a unitig with no predecessors. Hence, one unitig is extracted from the BBF instead of two.
- False branching: A false positive *k*-mer connects as a successor, respectively predecessor, to a true positive *k*-mer which already has a successor, respectively predecessor. Hence, three unitigs are extracted from the BBF instead of one.

An example of a cdBG containing the two types of errors is illustrated in Figure 3: K-mer ‘CCG’ creates a false branching and ‘ACT’ creates a false connection.

**Figure 3:**
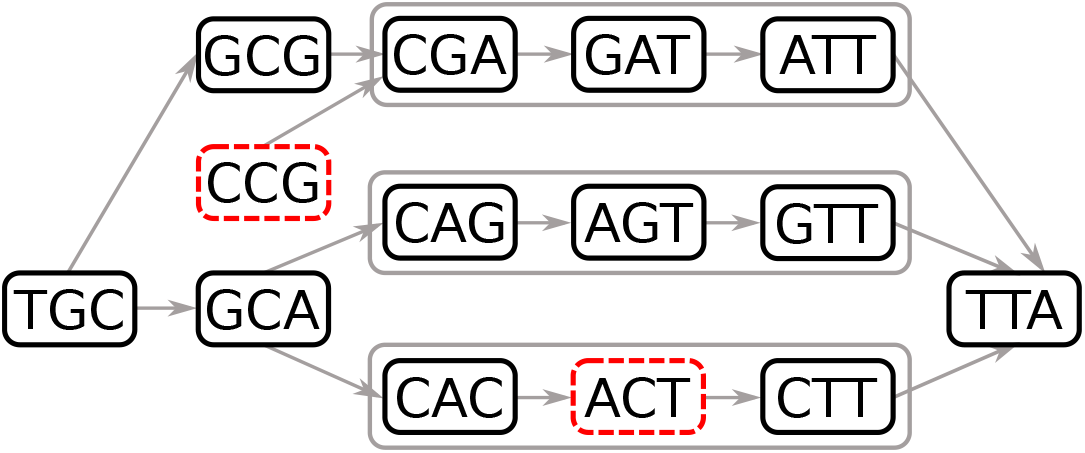
A compacted de Bruijn graph containing false positive 3-mers. Errors are represented in red dashed line vertices: K-mer ‘CCG’ creates a false branching and ‘ACT’ creates a false connection. K-mers that are compacted in a unitig are grouped in a grey line box.

In order to distinguish false positive from true positive *k*-mers, a counter is maintained on each *k*-mer of the unitigs and Algorithm 6 is modified to increment the counters of the *k*-mers occurring in the reads. Hence, false positive *k*-mers with no or one single occurrence are deleted from the graph. In the case of a false connection *k*-mer, deleting the *k*-mer splits a unitig. In case of a false branching, deleting the *k*-mer joins one or multiple unitigs.

#### 3.2.4 Ghost *k*-mers

The false positive rate of the BBF will affect the length of the unitigs extracted by Algorithm 5. Consider a unitig of length *k* + *η −* 1 in the true cdBG, consisting of *η k*-mers. For each internal *k*-mer, the algorithm makes 8 queries to the BBF, two of which will return true and 6 of which should return false. If the BBF has a false positive rate of *p*, the algorithm will advance to the next k-mer with probability (1 *− p*)^6^ *≈* 1 *−* 6*p* and stop prematurely with probability *≈* 6*p*. The number of *k*-mers in the extracted unitig will then be limited by *η* on one hand and a geometric distribution with probability 6*p*, whose expected value is 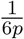. When *p* = 10^−3^ this would lead to an average unitig length of 167. While these errors are fixed with Algorithm 7, this leads to an increased memory usage. One way to increase the length would be to use more memory in the BBF which would reduce the false positive rate. However we observe that the most likely configuration is that a single false positive *k*-mer *x*′, adjacent to a real *k*-mer *x* in the unitig, causes a premature halt to the extraction of the true unitig. When *x*′ has no other neighbor in the BBF except for *x*, we call it a ghost *k*-mer, insert it into a hash table to keep track of it in case we observe it later but do not stop the extraction of the unitig. In the rare case that *x*′ turns out to belong to the true cdBG, we identify the unitig containing *x*′ and fix the mistake. The probability that we halt can now be approximated as 42*p*^2^, since this would require two adjacent false positive *k*-mers to occur in the BBF. The use of ghost *k*-mers greatly reduces fragmentation which improves memory usage and running time.

#### 3.2.5 Recurrent minimizers

Even in the case of a minimizer random ordering as described in Section 2, some minimizers are expected to occur more often in unitigs than others, due to indels occurring in homopolymer and tandem repeat sequences. Those minimizers are likely to increase the running time as their lists of tuples in the minimizer hash-table *M* will be much longer than for the other minimizers. We define a minimizer as *recurrent* if it occurs *t* times or more in the unitigs of the cdBG. In order to limit the impact of recurrent minimizers on the graph construction, list of tuples in *M* have a maximum length *t*. When a *k*-mer *x* and its corresponding minimizer *y* must be inserted into the cdBG data structure, the length of the list associated with *y* in *M* is verified first. If the length is greater or equals to *t*, *y* is a recurrent minimizer. In such case, a non-recurrent minimizer *y*′ > y is extracted from *x* and inserted into *M*. If *x* does not contain a non-recurrent minimizer *y*′, the recurrent minimizer *y* is inserted into *M* instead. Whenever *k*-mer *x* is searched, the list of tuples associated with its minimizer *y* is traversed and *x* is anchored on the instances of *y* in the unitigs of the graph until a match is found, as described in Algorithm 4. However, if no match is found for *x* and the list of tuples associated with *y* contains *t* or more tuples, the non-recurrent minimizer *y*′ is extracted from *x* and the search continues using minimizer *y*′.

##### Algorithm 7 Removal of False Positives

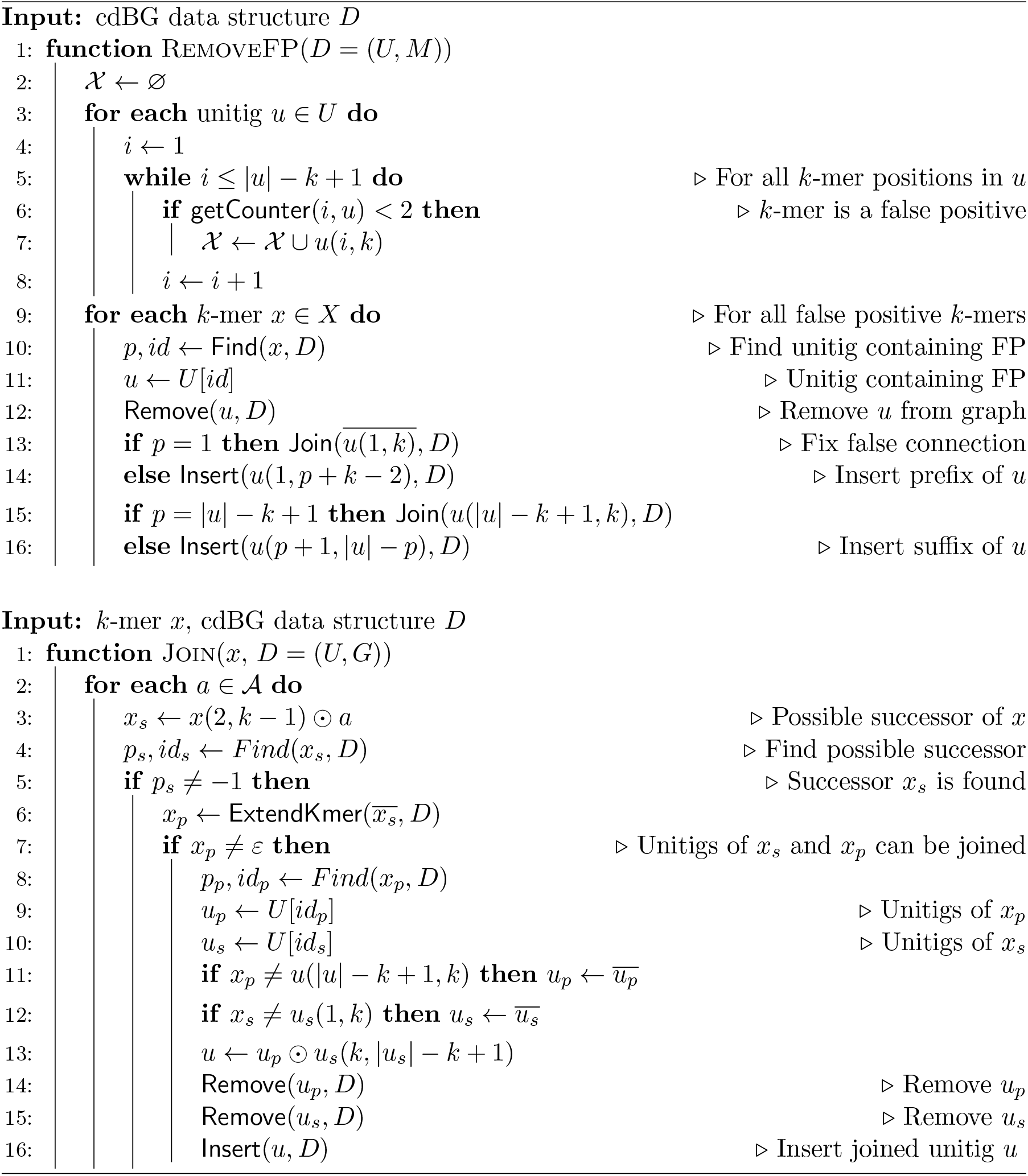

### 3.3 Coloring

We denote as *D*′ the data structure of a ccdBG: It is composed of a unitig array *U*, a minimizer hash-table *M*, an array *O* of color containers, an array *H* of hash functions and a hash table *K* of *k*-mers.

#### 3.3.1 Container representation

In Bifrost, a color is represented by an integer from 1 to *|C|*. A unitig *u* composed of *η* = *|u| − k* + 1 *k*-mers is associated with a binary matrix of size *η × |C|*: rows represent the different *k*-mer positions in *u* and columns represent the colors from *C*. A bit set at row 1 *≤ i ≤ η* and column 1 *≤ j ≤ |C|* indicates that *k*-mer *u*(*i, k*) occurs in dataset *j*. In order to limit the memory usage of colors, multiple compressed index are used to represent these binary matrices depending on their sparsity:

- A 64 bits word that can be either a tuple (position *i,* color *j)* or a binary matrix of size *η × |C| ≤* 62 (2 bits are reserved for the meta-data)
- A compressed bitmap adapted from a Roaring bitmap container (Chambi *et al.*, 2016). This compressed bitmap stores up to 65488 tuples (position *i,* color *j)* and uses a maximum of 8 KB of memory. This container has 3 representations of the tuples it indexes: bit vector, sorted list of tuples and run-length encoded list of sorted tuples. Compared to a Roaring bitmap, this compressed bitmap uses less memory for its meta-data and requires less cache-miss to access the tuples.
- A Roaring bitmap (Chambi *et al.*, 2016) to store more than 65488 tuples. Roaring bitmaps are SIMD accelerated and propose numerous functions to manipulate bitmaps such as set intersection and union.

Those representations have a logarithmic worst-case time look-up and insertion.

#### 3.3.2 Container indexing

Color containers can become substantially large and in order to avoid costly data transfer operations when the ccdBG data structure *D*′ is modified, color containers are not associated directly to unitigs in *D*′. Instead, a solution derived from the MPHF (Minimal Perfect Hash Function) library BBHash (Limasset *et al.*, 2017) is used to link unitigs of array *U* to color containers of array *O*. The benefit of such a method is that operations which affects only the structure of the graph do not move the color containers in memory. Algorithm 8 describes how color containers are associated to their respective unitigs.

## 4 Results

To compare the performance of Bifrost, we benchmarked against state-of-the-art software on publicly available dataset. We focus on two representative problems for which the de Bruijn graph has been employed, namely whole genome assembly of HTS short reads and pan-genome analysis of assembled genomes. All experiments were run of a server with an 16-core Intel Xeon E5-2650 processor and 256G of RAM. Running time was measured as wall clock time using the time command, peak memory was measured by ps. All programs that support multithreading were run with 16 threads.

### Algorithm 8 ccdBG Filters

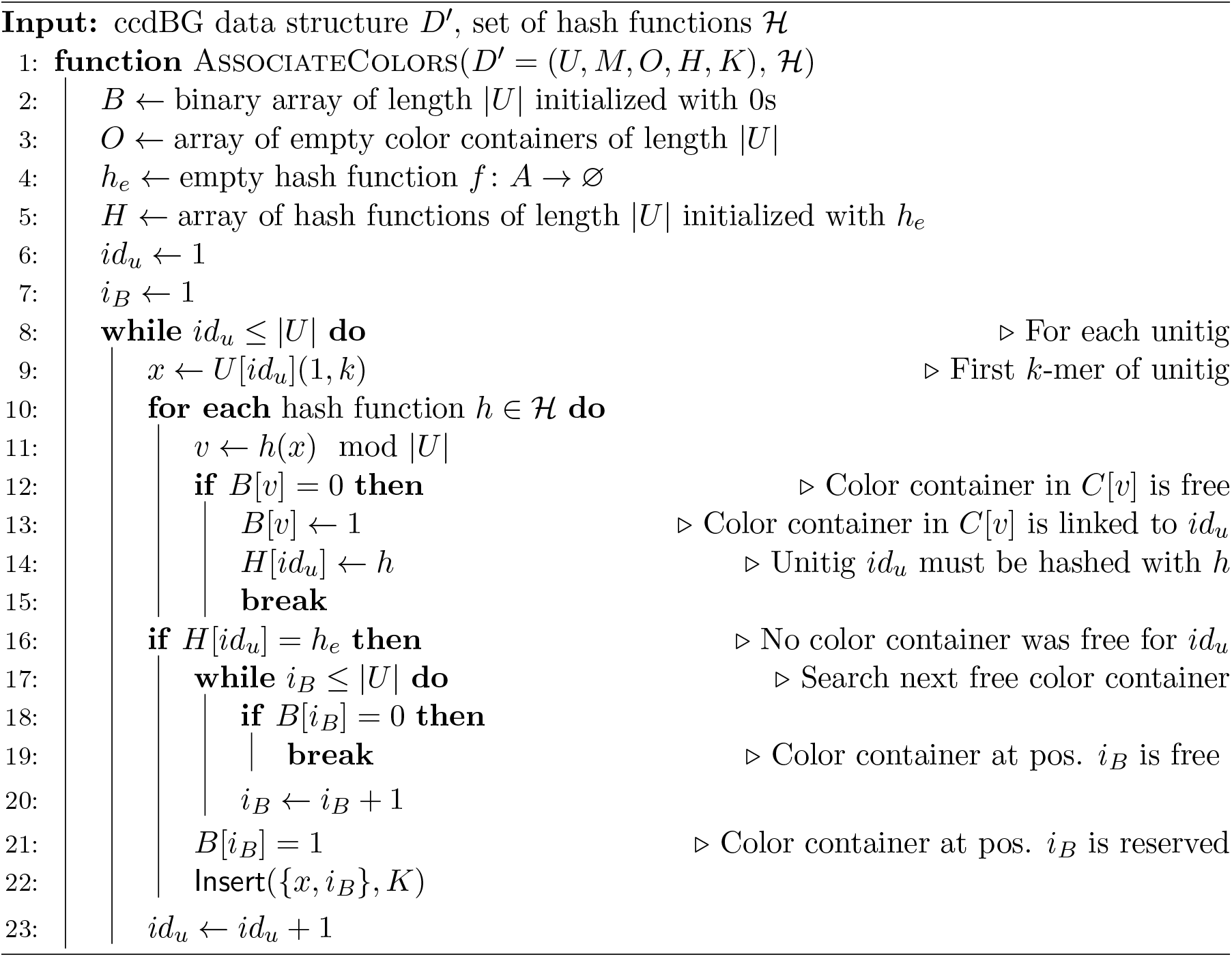

### 4.1 cDBG construction for Genome assembly

We constructed the compacted de Bruijn graph using a human genome short read dataset from the NA12878 sample from Genome In A Bottle consortium (Zook *et al.*, 2016), downsampled from 300-fold coverage to 30-fold coverage to reflect normal sequencing depth. The input is about 696 million 150 bp paired-end sequences with about 30x coverage. We compared Bifrost to the pipeline BCALM2 (Chikhi *et al.*, 2016) and Blight (Marchet *et al.*, 2019). BCALM2 is a disk-based compaction tool for short read data and assembled genomes while Blight is a non-dynamic data structure for indexing unitigs.

BCALM2 can be configured for different memory usage where a lower memory usage results in a longer running time. Hence, BCALM2 was configured in Table 1 with a maximum memory usage of 45 GB, similar to the Bifrost memory usage for the same dataset. We also ran BCALM2 with a 100 GB of memory, resulting in a running time of 4.91 hours and memory usage of 88 GB with no external disk used.

**Table 1:** Running time, memory usage, and external disk space used for de Bruijn graph building and indexing. Numbers in parenthesis show the respective use of BCALM2 and Blight: The running time was added while the maximum was taken for memory and disk.

|  | Running time (h) | RAM peak (GB) | Disk peak (GB) |
| --- | --- | --- | --- |
| Bifrost | 5.51 | **41.5** | **0** |
| BCALM2+Blight | **4.78** (3.16+1.61) | 46.6 (46.6,10.4) | 129 (129,1) |

### 4.2 Pan-genome analysis

We constructed colored and compacted de Bruijn graphs for a maximum of 117,913 assembled genomes of *Salmonella*. The input represents all publicly available *Salmonella* assemblies from the database Enterobase (Zhou *et al.*, 2019) as of August 2018. This is a 7.3x increase in the number of colors compared to the work of Muggli *et al.* (2019) who reported the construction for 16,000 *Salmonella* strains. We compared Bifrost to VARI-merge (Muggli *et al.*, 2019) as both tools can construct the colored de Bruijn graph and update it without reconstructing the graph entirely. The main differences between the two tools is that VARI-merge is mainly a disk-based method that produces a non-compacted colored de Bruijn graph. We only benchmarked VARI-merge as it is currently the state-of-the-art for colored de Bruijn graph construction. A comparison of VARI-merge to other colored de Bruijn graph construction tools is given in (Muggli *et al.*, 2019). Results are given in Table 2 for a variable number of strains. Note that the reported VARI-merge time includes the time spent by KMC2 (Deorowicz *et al.*, 2015) to compute the *k*-mers required in input of VARI-merge.

**Table 2:** Running time, memory usage, and external disk usage for constructing the colored de Bruijn graphs of an increasing number of *Salmonella* strains. N/A indicates the result is unavailable.

|  | Number of strains | Running time (h) | RAM peak (GB) | Disk peak (GB) |
| --- | --- | --- | --- | --- |
| Bifrost | 100 | **0.016** | **0.16** | **0** |
| VARI-merge |  | 0.33 | 5.1 | 17 |
| Bifrost | 400 | **0.05** | **0.29** | **0** |
| VARI-merge |  | 1.016 | 15.4 | 51 |
| Bifrost | 1600 | **0.38** | **2.4** | **0** |
| VARI-merge |  | 4.86 | 56.9 | 228 |
| Bifrost | 4000 | **1.66** | **3.7** | **0** |
| VARI-merge |  | 12.35 | 138 | 449 |
| Bifrost | 117913 | **93.35** | **102.74** | **0** |
| VARI-merge |  | N/A | N/A | N/A |

In Muggli *et al.* (2019), the authors process 16,000 strains in batches of 4,000, merging the batches to produce a colored de Bruijn graph of all strains. This required 254Gb of memory and 2.34Tb of external disk, with a total running time of 69 hours. In comparison, Bifrost processed 117,913 strains using about 103Gb of memory, no external disk usage and a total running time of 93.35 hours. While the running time is not directly comparable across different machines due to different processors, this is in line with Bifrost being about eight times faster than VARI-merge.

## 5 Discussion

We present Bifrost, a method for constructing compacted de Bruijn graphs, both regular and colored, with minimal memory requirements. Bifrost is competitive with the state-of-the-art de Bruijn graph construction method BCALM2 and the unitig indexing tool Blight with the advantage that Bifrost is dynamic. For colored de Bruijn graphs, Bifrost is about eight times faster than VARI-merge and uses about 20 times less memory with no external disk. The query capabilities of Bifrost are both for identifying colors for a given *k*-mer as well as navigating the de Bruijn graph. The software was developed with the intention of being usable as a tool or a library wherever large de Bruijn graphs are needed with minimal external dependencies.

## Acknowledgements

The authors would like to thank Nina Luhmann, Birte Kehr and Thomas Krannich for their helpful feedback during the development of the software and Trausti Sæmundsson for work on an early draft of the software.

